# Advancing ecological assessment: The integration of eDNA metabarcoding into an estuarine fish index

**DOI:** 10.1101/2025.03.31.645977

**Authors:** Mukesh Bhendarkar, Oriol Canals, Carlos Jurado, Iñaki Mendibil, Ainhize Uriarte, Angel Borja, Naiara Rodríguez-Ezpeleta

## Abstract

In the face of increasing anthropogenic pressures on estuarine ecosystems, methods to efficiently and reliably assess their ecological status are essential. This study explores the integration of environmental DNA analysis into the AZTI’s Fish Index (AFI) to assess ecological status of estuarine ecosystems. Surface water eDNA sampling and bottom trawl surveys were performed across multiple estuaries in the Basque Country, Spain, and resulting species data were used to calculate AFI scores. eDNA metabarcoding consistently detected higher fish species richness than bottom trawling, while the latter remained more effective at capturing demersal species. In general, ecological classifications from eDNA- and bottom trawl derived data displayed low concordance, largely due to differing species assemblages and metric contributions. These results emphasize the respective strengths and weaknesses of each methodology and the necessity for method-specific calibration. Considering that the AFI is calibrated using bottom trawl data, its direct application to eDNA-derived species lists may lead to some inconsistencies. This study underscores the critical necessity to establish eDNA-specific reference conditions and to recalibrate index thresholds accordingly. While eDNA approach may not entirely replace traditional methods, its scalability, sensitivity, and minimal ecological disturbances establish it as an essential complementary application within estuarine assessment programs. This research strongly supports the urgent advancement of eDNA-based indices and the critical enhancement of reference conditions for their effective incorporation into ecological assessment frameworks under the Water Framework Directive.

## Introduction

Estuaries, functioning as transitional zones between freshwater and marine environments, are amongst the most productive ecosystems on Earth, offering vital ecosystem services to nearby communities (Adey, 2024; Barbier et al., 2024). The ecological importance of estuaries lies not only in their biodiversity but also in their ability to regulate water quality and provide habitats for various species. However, anthropogenic activities and environmental stressors increasingly threaten these ecosystems, leading to significant reductions in ecosystem services (Allen et al., 2023; Elliott & Kennish, 2024; Jennerjahn & Mitchell, 2013). Additionally, climate change exacerbates these threats, raising concerns on the sustainability of estuarine supply of services (Siemes et al., 2024).

In this context, assessing the ecological status of estuaries is a priority, and various assessment methods have been established under the European Water Framework Directive (WFD) for different biological elements (Birk et al., 2012) such as phytoplankton, benthic flora, benthic invertebrates and fish. Each of these methods varies between countries due to differences in the response of indicator species to relevant stressors, local species composition, sampling methods and available taxonomic resolution based on regional knowledge of flora and fauna (Birk et al., 2012; Borja et al., 2013). One fundamental element of these methods is the use of a “reference condition” which is specific to each aquatic ecosystem, from which Ecological Quality Ratio (EQR) is derived by assessing the deviation of the calculated value from this reference (European Commission, 2000; Van De Bund & Solimini, 2007). However, all of them have been harmonized through intercalibration among countries using them (European Commission, 2024).

Several fish-based indices have been proposed in Europe (Pérez-Domínguez et al., 2012) and globally (Cabral et al., 2022), such as the AZTI’s Fish Index (AFI), developed primarily for evaluating ecological quality in the Basque Country estuaries, in Spain (Borja et al., 2004; Uriarte & Borja, 2009). Nowadays, AFI is one of the most widely used indices for assessing the ecological quality and biotic integrity of transitional waters (Souza & Vianna, 2020). It assesses EQR using composition and abundance data from bottom trawl surveys. However, sampling in estuarine environments presents significant logistical and technical challenges associated with physical capture of species (Testa et al., 2017). Additionally, different capture methodologies exhibit biases towards certain faunal behaviour, which culminates in incomplete evaluations of the structure of estuarine species compositions (Franco et al., 2012). For instance, bottom trawling predominantly targets benthic-demersal species, consequently restricting inferences to the pelagic zone while misidentification of juvenile and cryptic species complicates capture-based methods (Kirsch et al., 2018). These methods require extensive fieldwork, expert taxonomic knowledge, and are invasive, potentially disturbing sensitive ecosystems and organisms they aim to conserve (Bortolus, 2008; Stribling et al., 2008). Therefore, advancing faster, more precise, and non-invasive methods to assess the ecological status of estuaries is essential.

Given these limitations, environmental DNA (eDNA) metabarcoding has emerged as a promising tool for aquatic biodiversity assessments. By analysing genetic material shed by organisms into the environment, eDNA enables the detection of species without the need for physical capture. Although applicable to a broad range of taxa, recent applications have focused on its potential for monitoring fish communities in transitional waters (Beng & Corlett, 2020; Nagarajan et al., 2022). It has already been use at the national biomonitoring efforts, especially for Special Areas of Conservation (SAC) under the EU Habitats Directive (Gallagher et al., 2020; Jacobsen et al., 2023), reflecting its growing importance for integration in biological assessment of aquatic ecosystems (Pawlowski et al., 2018). Thus, it stands out as a non-invasive option to complement and potentially replace traditional methods, while offering a new paradigm for ecological assessment that is consistent with the principles of WFD (Borja et al., 2024; Hering et al., 2018).

In this study, we explore the integration of eDNA metabarcoding into the AZTI’s Fish Index (AFI) to assess its potential for evaluating the ecological status of estuarine ecosystems. Specifically, we compare eDNA-based assessments with those derived from traditional bottom trawl surveys to examine their relative performance, agreement in ecological classification, and suitability for regional monitoring frameworks. We hypothesize that eDNA can provide results broadly consistent with trawl-based assessments, while also offering added value through improved species detection and non-invasive sampling. By identifying key areas of convergence and divergence between the methods, this study aims to inform the advancement of eDNA-based tools for future ecological assessment and their potential role in supporting Water Framework Directive implementation.

## Material and Methods

### Study area and sample collection

The study was carried out along the Basque estuaries of Spain during Autumn of 2017, 2018 and 2019, focusing on the estuaries of rivers Artibai, Barbadun, Bidasoa, Butroe, Deba, Lea, Nerbioi, Oiartzun, Oka, Urola and Urumea, as part of the Basque Monitoring Network, from the Basque Water Agency (URA). eDNA sampling in surface waters and bottom trawling were carried out in three sections (inner, middle and outer) of each estuary **(Figure 1; Table S1)**.

**Figure 1.**
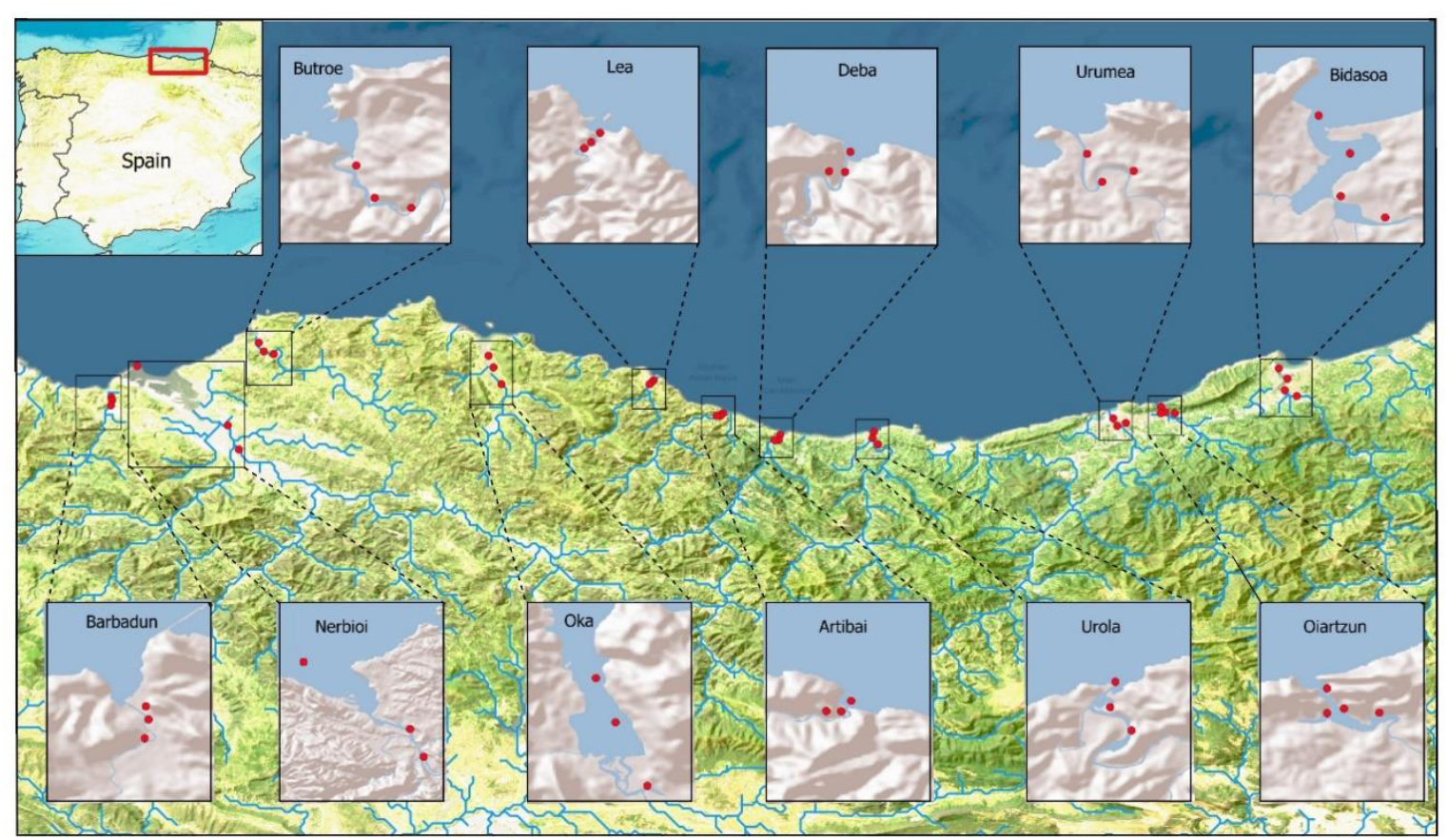
Map of the study area and sampling sites. Red dots represent the specific sampling locations in each section.

Surface water samples were collected prior to bottom trawling operations. Samples were collected in 5 litre plastic containers and transported to the laboratory under cold and dark conditions without exposure to trawl catches. In the laboratory, the water samples were pre-filtered through a 60 μm mesh-size net and then filtered through 0.45 μm pore size Sterivex filters (Millipore) using a peristaltic pump. The filters were stored in plastic bags at −80 °C until further analysis. After filtration, and before the next sampling, the containers and filtration system were thoroughly decontaminated by washing extensively with 10% bleach, rinsed with Milli-Q®water, dried, and stored at room temperature.

The demersal community (fish and crustaceans) was sampled using a beam trawl 1.5 m wide and 300 mm high, with a mesh size of 40 mm at the wings and 8 mm at the cod-end. The net was trawled at a speed of 1.5-2 knots for 10 min during high or rising tide, with three replicates. The collected specimens were placed in a box containing water and anaesthetic. This was performed to facilitate morphological identification and fish measurements. All procedures involving fish were carried ensure minimal stress and harm. Taxonomic identification was performed at the lowest possible taxonomic level (i.e. species), with individual counts recorded for each sampling location. Fish were released back to water immediately, except for those specimens (either fish or crustaceans) that could not be identified, which were preserved in ethanol.

### DNA extraction and amplicon library preparation

DNA was extracted using the DNeasy® blood and tissue kit (Qiagen) following at modified protocol for DNA extraction from Sterivex filters without preservation buffer by Spens et al. (2017). DNA concentration was measured with at Quant-iT dsDNA HS assay kit using a Qubit® 2.0 Fluorometer (Life Technologies, California, USA). Library preparation was performed following a modified version of the Tagsteady protocol (Carøe & Bohmann, 2020). Each sample was amplified in triplicate using a different set of tagged primers including 6 bases for tag, one or two degenerated bases (Ns) and the *teleo* primer pair (forward: 5’ACACCGCCCGTCACTCT3’; reverse: 5’CTTCCGGTACACTTACCATG3’) (Valentini et al., 2016), targeting a 60-70 bp long region of the mitochondrial 12S rRNA gene. PCR mixtures were prepared under the hood in the pre-PCR laboratory using dedicated micropipettes and disposable plastic ware that were previously decontaminated under UV light, and all post-amplification steps were carried out in the post-PCR laboratory. PCR amplifications were done in a final volume of 15 μl including 7.5 μl of 2X Phusion Master Mix (ThermoScientific, Massachusetts, USA), 0.3 μl of each amplification primer (final concentration of 0.2 μM), 3 μl of teleo_blk (final concentration of 2 μM) (Valentini et al., 2016), 2.5 μl of MilliQ water, and 2 μl of 10 ng/μl template DNA. The thermocycling profile for PCR amplification included 3 min at 98 °C; 40 cycles of 10 s at 98 °C, 30 s at 55 °C, 45 s at 72 °C, and finally 10 min at 72 °C. Triplicates of PCR products were pooled and differently tagged PCR products were pooled at approximately equimolar ratios based on gel band strength. The mixes were then purified using AMPure XP beads (Beckman Coulter, California, USA) following the manufacturer’s instructions, and Illumina’s compatible adapters were ligated to each mix as in Carøe and Bohmann (2020). Libraries were quantified using the Quant-iT dsDNA HS assay kit using a Qubit® 2.0 Fluorometer (Life Technologies, California, USA) and pooled in the same quantity for sequencing on the Illumina NovaSeq platform (Illumina, California, USA) with a 2 × 150 paired end protocol. One positive control (DNA extracted from cat fur) and one to two PCR negative controls were added to each mix to detect possible cross contaminations. Additionally, sequences from non-used adapter combinations were retrieved to account for index hopping.

### Read pre-processing and taxonomic assignment

Raw reads were demultiplexed using cutadapt (Martin, 2011) based on tags not allowing any error (−e 0), and quality was verified using FASTQC (Andrews, 2010). Paired reads were merged using Pear (Zhang et al., 2014) with a minimum overlap of 30 nucleotides, and merged reads with an average quality lower than 25 Phred score were removed using Trimmomatic (Bolger et al., 2014). The degenerated bases added during library preparation and the forward and reverse primers were removed using cutadapt (Martin, 2011) allowing an error rate of -e 0.2. Since in the Tagsteady protocol both forward and reverse-oriented reads occur in both the R1 and R2 files, primer trimming was carried out in parallel for forward and reverse reads, and reads were joined later. Reads that: i) did not cover the teleo region, ii) were shorter than 50 nucleotides, and iii) contained ambiguous positions, were removed using mothur (Schloss et al., 2009), as well as potential chimeras, which were detected using the UCHIME algorithm (Edgar et al., 2011). Taxonomic assignment of unique reads was performed according to the naïve Bayesian classifier method from Wang et al. (2007) implemented in mothur using a 70% confidence cutoff threshold, against a curated, local reference database built as described by Claver et al. (2023), which includes 924 fish species expected in the Northeast Atlantic and Mediterranean areas. Only those sequences identified at the genus or species level were considered for subsequent analyses. Taxa detected with less than 0.1% of reads in a sample were excluded to reduce the potential for spurious sequences (Drake et al., 2021; Skelton et al., 2023).

### AFI calculations

The AFI was calculated using nine distinct metrics: species richness, percentage of pollution indicator species, introduced species, fish health, flat fish presence, omnivorous species, piscivorous species, number of resident species, and percentage of resident species (Borja et al., 2004; Uriarte & Borja, 2009). For each species listed in the AFI framework **(Table S2)**, a value of 0 or 1 was assigned across seven ecological indicator categories, depending on the species’ characteristics. These values were then summed for all species detected in a sample to generate the AFI score. The species included in the AFI list were based on observations from Basque estuaries (via studies by URA, the Consorcio de Aguas Bilbao-Vizcaya, and Gipuzkoa Council) as well as other European transitional waters. The AFI was computed by correlating the indicator values into scores of 1,3, or 5 and the total of all scores resulted in the final quality classification **(Table S3)**. Considering that eDNA metabarcoding does not enable assessing fish health, the fish health indicator was uniformly assigned a maximum score of 5 in both the trawl and eDNA based data. Quality classification consists of five scores: high from 39 to 45; good from 31 to 38, moderate from 24 to 30, poor from 17 to 23 and bad from 9 to 16 (Borja et al., 2004). These scores serve as indicators of an EQR between 0 to 1, where a score of 9 corresponds to 0 (bad quality) and a score of 45 corresponds to 1 (high quality).

For the AFI based on the trawl survey, two versions were calculated. The first version included all trawled fish species as well as crustaceans, which were included in the AFI list to represent the demersal component and enhance classification due to the small number of resident species in the Basque estuaries (Borja et al., 2004). The second version included only fish species, and no crustaceans. Similarly, for the eDNA-based AFI, two versions were calculated. The first version included all fish species detected through eDNA metabarcoding that are part of the AFI list **(Table S2)**. The second version considered only those species that occurred at least once in the trawling efforts conducted over 18 years within the estuaries of the study area (compiled as ‘historical database’ **Table S4**), subsequent to the work of Borja et al. (2025). A genus-level approach (Little, 2011) was used to account for potential misidentifications due to incompleteness of the reference database (Claver et al., 2023) or limited taxonomic resolution of the *teleo* barcode (Collins et al., 2019). This grouping at the genus level was done for species of the same genus exhibiting same scores only **(Table S5)**. For example, we merged the *Diplodus* species showing the same scoring (*Diplodus vulgaris, D. sargus* and *D. cervinus*) but retained *D. annularis* as a separate taxon **(Table S6)**.

### Statistical analysis

All statistical computations and visualizations were executed utilizing R environment version 4.0.4 (R Core Team, 2021) with the following packages: tidyverse, dplyr, VennDiagram, ggplot2, ggpubr, ggrepel, gridExtra, and cowplot. Pair and boxplots were used to compare the performance of different AFI score derived from eDNA and bottom trawl data. Cohen’s Kappa (κ) was performed to determine the inter-rater of agreement between the AFI results obtained for obtained across different versions. The level of agreement is categorized as described by Monserud and Leemans (1992); no agreement (κ ≤ 0.05), very poor (0.05 ≤ κ ≤ 0.20), poor (0.21 ≤ κ ≤ 0.40), fair (0.41 ≤ κ ≤ 0.55), good (0.55 ≤ κ ≤ 0.70), very good (0.71 ≤ κ ≤ 0.85), Excellent (0.85 ≤ κ ≤ 0.99), and perfect (0.99 ≤ κ ≤ 1.00). In addition, Kruskal–Wallis rank sum tests were used to assess differences in species richness detected by each method. Subsequently, post hoc analyses were conducted using Dunn’s test. Student’s t-test was conducted to assess the differences in AFI metrics (richness, pollution indicators, flatfish, omnivorous, piscivorous, and resident species) between the eDNA-AFI and trawl survey. The Relative Contribution Ratio (RCR) was calculated to determine the proportional contribution of each ecological metric to the AFI scores (Luo et al., 2015). For each method, RCR was computed as the percentage contribution of a given metric to the total AFI score. All of these analysis custom scripts and output files utilized for analysis are deposited in a public repository at https://github.com/mukeshbhendarkar/eDNA-AFI.

## Results

### Sampling efficiency of eDNA and bottom trawling

All samples together, eDNA metabarcoding yielded a total of 16,496,674 reads corresponding to 73 taxa (60 taxa taxonomically assigned to the species level and 13 at the genus level) from 30 orders and 37 families **(Table S7)**. Mullets were the dominant group and accounted for approximately 46% of the reads, with *Chelon ramada* (35.1%) and *Chelon labrosus* (7.7%) being the most abundant. Other frequently detected species with significant proportion of reads included *Sardina pilchardus* (8.8%), *Dicentrarchus labrax* (6.8%), and *Solea solea* (6.7%). On the other hand, the trawling survey captured 310 individual fish classified into 23 species or genus (1 specimen was identified to family level and two specimens remained unclassified). Gobies were the dominant group in the catches, with *Pomatochistus* sp. and *Gobius niger* being the leading species, representing 26% and 22% of total catch, respectively. Other abundant captures were the flat fish *Solea solea* (20%), and the seabream *Diplodus sargus* (14%).

As expected, eDNA metabarcoding consistently detected significantly higher fish species richness compared to the traditional trawling approach (p < 0.05) in all sampling sites except in three of them: AME (Barbadun), AOKI (Oka) and OIAE (Oiartzun) **(Figure S1)**. A total of 43 fish species identified through the eDNA survey are listed in the AFI database of which 24 were not recorded in the trawl-based historical database **(Figure 2)**.

**Figure 2.**
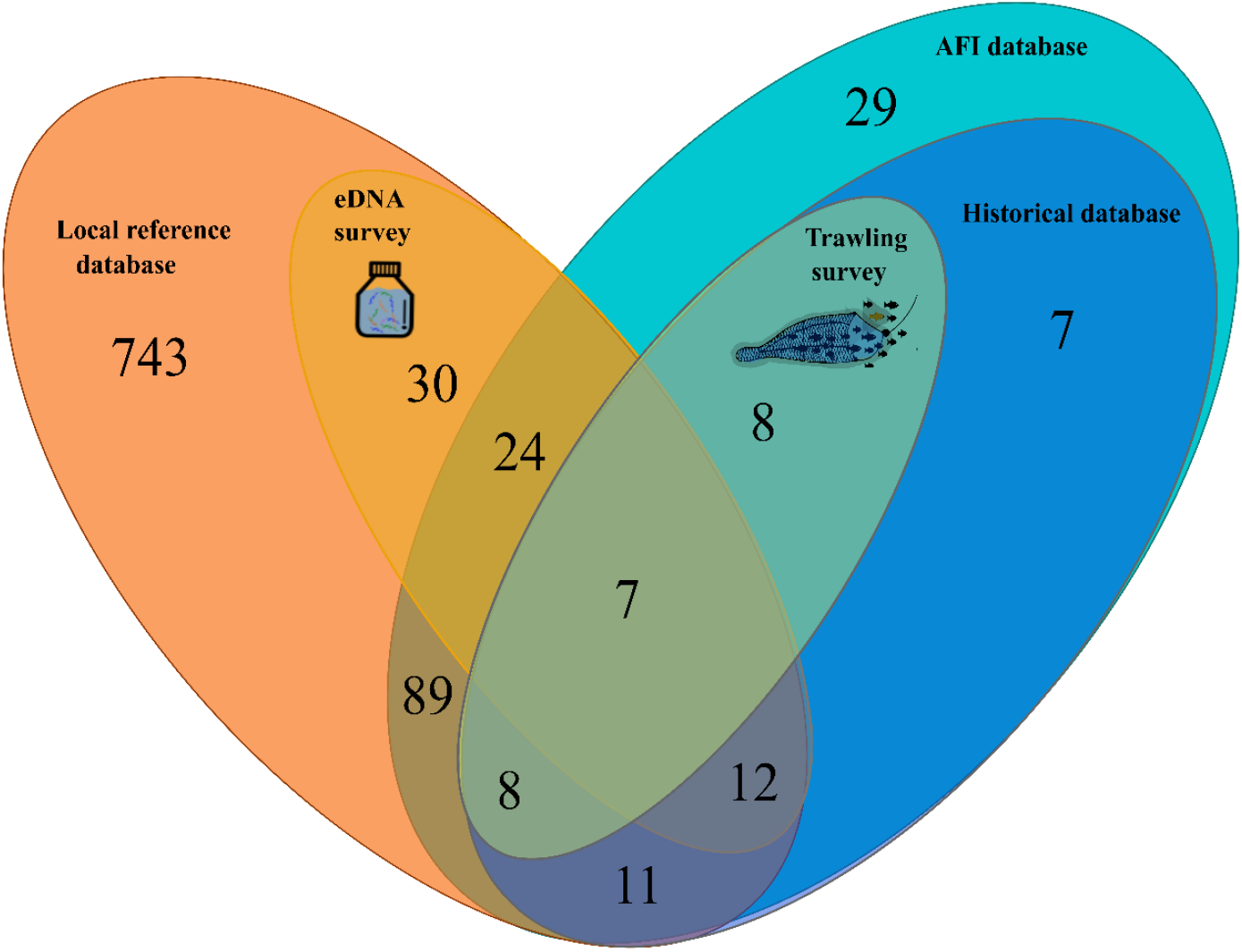
Venn diagram showing the number of species obtained in the eDNA metabarcoding (orange) and bottom trawling (green) surveys, with each method supported by a comprehensive reference database. The trawling survey’s reference includes historical trawl catch data from 2002-2019, while the eDNA survey utilizes a local sequence database from the Northeast Atlantic. It is important to note that the trawling reference database excludes species that remain unclassified up to the genus and/or species level.

Only seven species were recorded using both methods, yet they accounted for 87% of the trawl caught individuals and 24% of the eDNA sequence reads. The trawling survey captured 16 fish taxa that were not detected through eDNA metabarcoding. Of these, eight taxa were absent in the reference database, while the remaining eight, despite being in the reference database, were not observed in the eDNA metabarcoding. Conversely, eDNA metabarcoding exclusively detected 12 taxa within the historical database and 30 additional taxa not listed in the AFI database, many of which accounted for less than 1% of total reads **(Figure 2; Table S7)**.

### Assessing trawl and eDNA-based estuarine ecological status

AFI scores were calculated using both eDNA metabarcoding- and bottom trawl-derived data with and without the inclusion of crustaceans. For eDNA metabarcoding, the analysis accounted for 35 different taxa according to the AFI list and 17 different taxa based on the historical database (genus-based). The bottom trawling based AFI computations included 25 fish taxa, in addition to 13 crustacean species. Disparities in ecological status classifications between these methodologies were observed. For instance, eDNA-based assessments categorized two sites as having a ‘high’ ecological status, whereas trawl-based evaluations classified these as ‘good.’ Conversely, four sites classified ‘high’ by trawl assessments ranged from ‘good’ to ‘poor’ based on eDNA-derived data **(Figure 3)**. Notably, when the AFI calculations were restricted to fish catch data, excluding crustaceans, the majority of sampling sites resulted in ‘moderate’ and ‘poor’ ecological status, with the exception of sampling site ANE, which indicated a high status, and sites ABM, OIAE, OIAI2, and UROM, which exhibited good status **(Figure 3b)**. Cohen’s kappa values provided insight into methodological agreement **(Figure 3a)**. The two eDNA-based AFI computations exhibited near-perfect concordance (κ = 0.90), reflecting consistent results across the two datasets. In contrast, the two trawl-based AFI methods (with and without crustaceans) showed minimal agreement (κ = 0.07), indicating variability in ecological status assessments when crustaceans were considered. Crucially, no agreement was observed between eDNA- and trawl-derived AFI values, with kappa values ranging from −0.07 to −0.53 **(Figure 3b)**.

**Figure 3.**
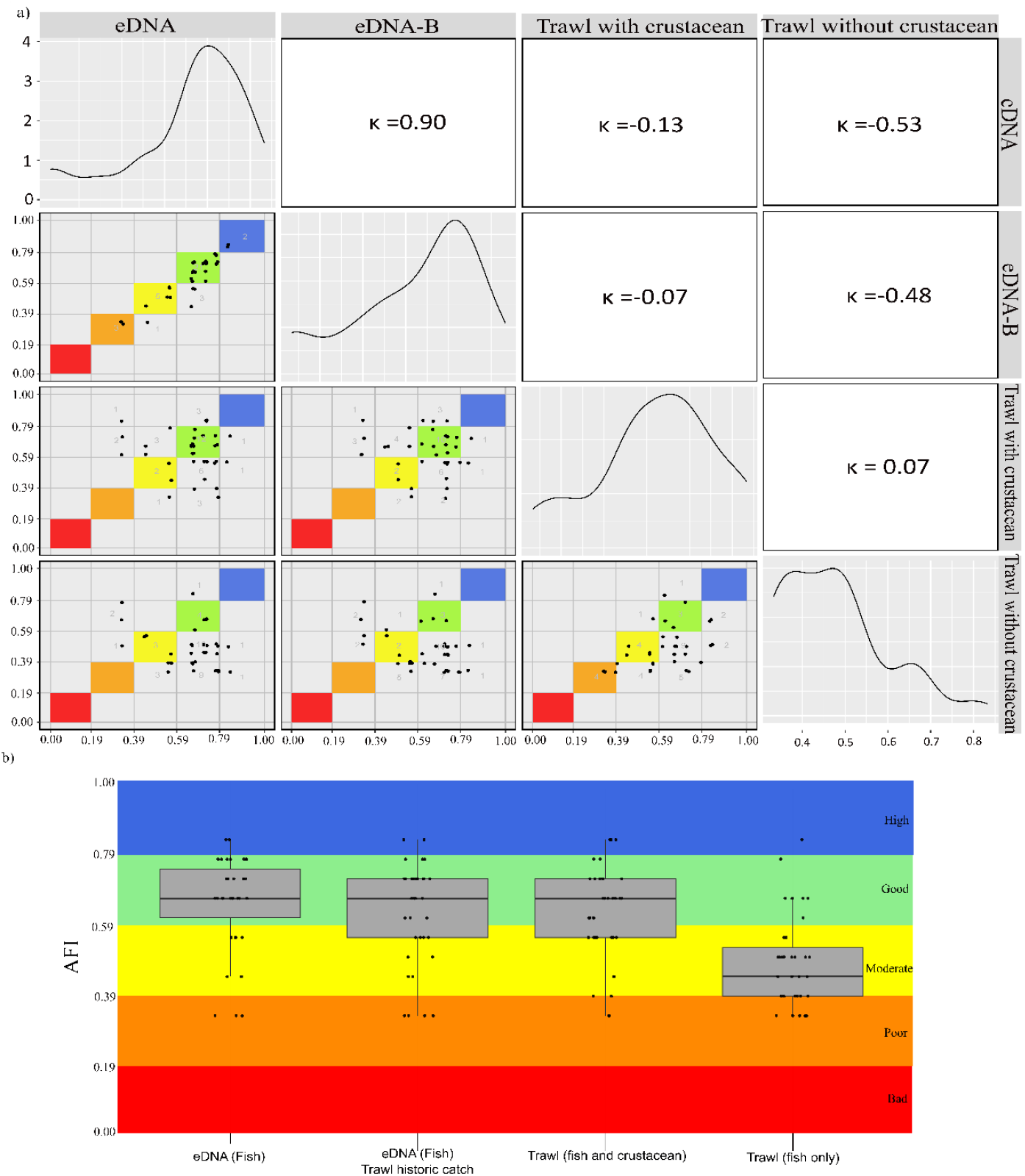
Comparison of AZTI’s Fish Index (AFI) scores across different approaches; eDNA (considering AFI listed species), eDNA-B (considering historical database species), Trawl with crustacean, and Trawl only fish (without crustaceans). a) the level of inter-rater concordance between eDNA metabarcoding and bottom trawl-derived data, with all versions. Cohen’s kappa (κ) values indicate the level of agreement. b) AFI scores across various sampling stations, with bars representing different computation approaches. Scores range from 0 (‘Bad’ status) to 1 (‘High’ status), with a colour gradient indicating ecological status. Some data points overlap due to identical AFI scores at different sites. Jitter (width = 0.01, height = 0.01) was applied to reduce overlap.

Additionally, significant differences were observed in some ecological metrics between methods (Student’s *t*-test, *p* < 0.05, **Figure 4**), namely species richness, pollution indicator species, proportions of flatfish, piscivorous species, and estuarine residents. The RCR analysis further revealed the influence of individual metrics on AFI scores. For eDNA-based assessments, omnivorous species contributed the most, accounting for 52.8% to 56.7% of the AFI score. Piscivorous species and resident species were secondary contributors, with 17.9% to 16.5% and 12.2% to 13.5%, respectively. In trawl-based assessments, contributions were more evenly distributed. Resident species had the highest impact (43.7% in fish-only analyses), followed by flatfish (28 %) and omnivorous species (17.6%).

**Figure 4.**
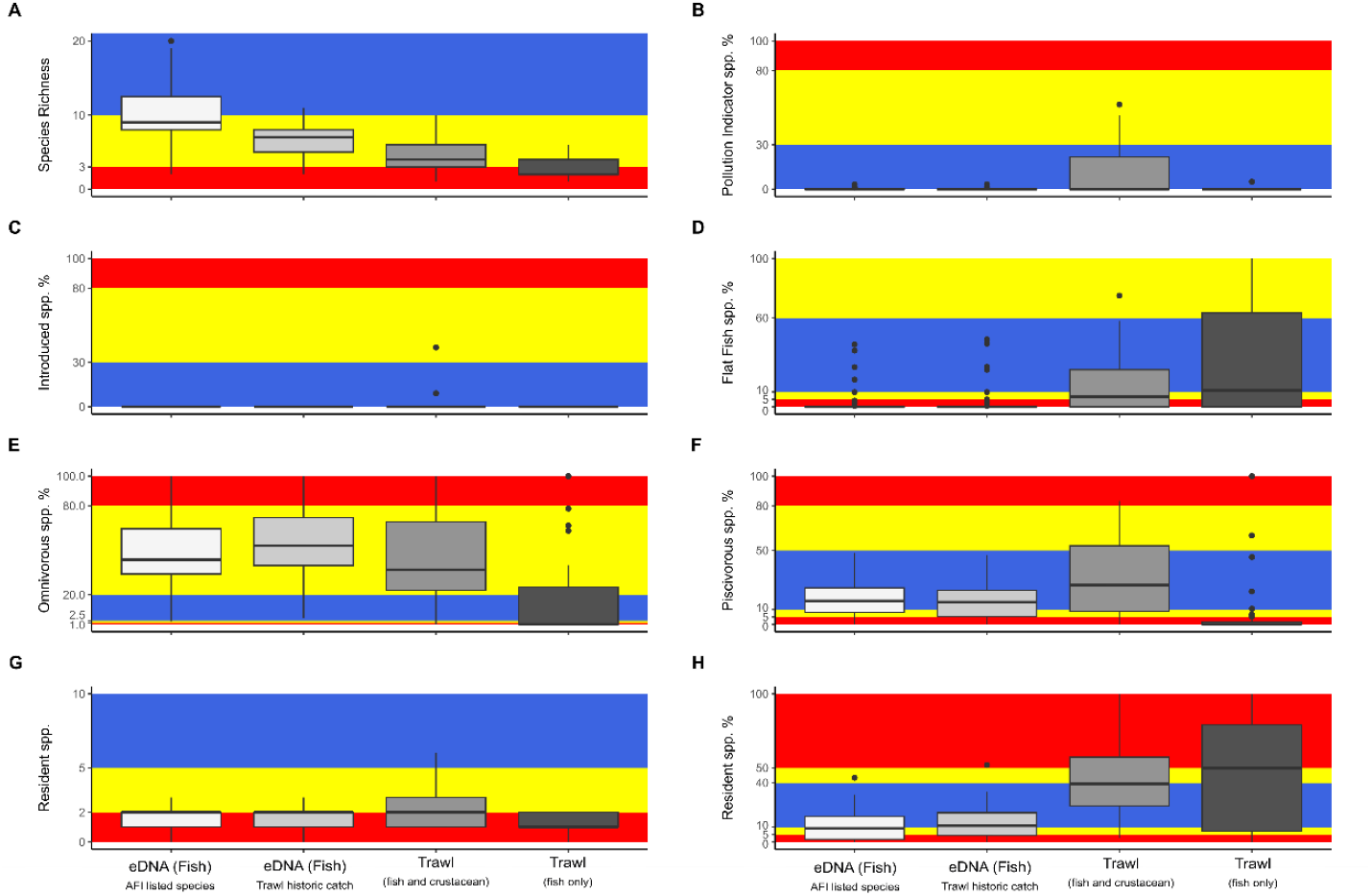
Influence of ecological indicators on AZTI’s Fish Index (AFI) score: (A) Species Richness, (B) Pollution Indicator Species (%), (C) Introduced Species (%), (D) Flat Fish Species (%), (E) Omnivorous Species (%), (F) Piscivorous Species (%), (G) Number of Resident Species, and (H) Resident Species The scores assigned indicate the overall status (see Table 1), with blue representing a score of 5, yellow a score of 3, and red a score of 1.

## Discussion

### Methodological contrasts

This study demonstrates the distinct capabilities of eDNA metabarcoding and bottom trawling in capturing estuarine fish assemblages, driven by their inherent methodological differences. eDNA metabarcoding detected over three times more fish species than bottom trawling, emphasizing its broader taxonomic reach. These findings align with previous studies demonstrating that eDNA consistently detects a higher number of fish species compared to conventional methods in estuaries (Gibson et al., 2024; Gibson et al., 2023; Saunders et al., 2024; Zou et al., 2020). However, this difference should not be interpreted as an inherent superiority of one method over the other, but rather as a reflection of their fundamentally differences in sampling approaches: eDNA was collected from the surface water during high tide and mainly detected pelagic and water-column-associated species, while bottom trawling primarily captured bottom-dwelling fish. In addition, eDNA detects genetic material present in the water column, allowing for the identification of species that may not be physically present at the time of sampling (Jerde, 2021), in contrast to bottom trawling, which is based on physical captures.

Among the identified species, only seven were common to both methods. These seven taxa, all demersal, were among the most abundant fish in bottom trawl catches (representing 87% of the total catch), yet they accounted for only 24% of total eDNA reads. Since eDNA concentration in the water is generally assumed to correlate with biomass (Nakagawa et al., 2022), their detection through eDNA metabarcoding suggests that highly abundant demersal species are more amenable to be detected in eDNA surface water samples than less abundant ones, which might remain undetected. Despite eDNA metabarcoding capturing a higher number of species overall, it still failed to detect 16 taxa captured by trawling, predominantly demersal (including flatfish) species. Among these, 8 species lacked reference sequences in publicly available genetic databases, preventing its detection, which reinforces the dependence of eDNA metabarcoding on comprehensive and regionally relevant reference databases for accurate species identification (Bhendarkar & Rodriguez-Ezpeleta, 2024; Claver et al., 2023; Marques et al., 2021). The remaining 8 species, despite being present in the reference database used for taxonomic classification, were not detected in eDNA samples likely due to their low abundance (and therefore expected low DNA concentration in the water) and/or limited vertical transport of their genetic material to surface water, suggesting the need of incorporating bottom water layers in eDNA sampling strategy to improve demersal species detection efficiency.

On the other hand, eDNA metabarcoding detected 12 species not captured by bottom trawling but previously recorded in the historical database, demonstrating its potential to detect species that may evade physical capture due to their behaviour, rarity, or habitat preference. These results align with previous studies suggesting that bottom trawling, while effective for demersal species, may overlook species better detected through eDNA (Afzali et al., 2021; Ip et al., 2024; Zhou et al., 2022). An additional 24 taxa previously unrecorded in the historical database were also detected through genetic means, demonstrating eDNA’s capacity to reveal potentially overlooked taxa. For instance, the most abundant species detected via eDNA metabarcoding—*Chelon ramada* and *Sardina pilchardus*—were not captured by bottom trawl and have not been previously documented in the historical database. The absence of *C. ramada* in bottom trawl surveys may be due to challenges associated with identifying mugilids through morphological traits, as they have historically been classified only to the family level in trawl records from Basque estuaries. Similarly, the absence of *S. pilchardus* in bottom trawl catches is likely attributable to its pelagic nature and ability to avoid bottom trawling gear (Haugland & Misund, 2011). While eDNA metabarcoding improves biodiversity detection, it also introduces uncertainties regarding the origins and ecological relevance of detected eDNA signals. One of the primary challenges in interpreting eDNA-based detections is the possible influence of eDNA transport mechanisms, particularly in estuarine environments characterized by complex hydrodynamics (McCartin et al., 2024). Thus, some species detections may result from eDNA fragments transported from upstream freshwater sources or introduced through tidal movements from the marine environment, rather than indicating actual species presence at the sampling location (Jeunen et al., 2019; Polanco et al., 2021). Additionally, contamination from anthropogenic sources such as sewage effluents or food waste could introduce foreign DNA, potentially inflating species richness estimates (Inoue et al., 2023). These factors highlight the need for refined analytical approaches to differentiate resident species from transient or external DNA sources, particularly in dynamic coastal ecosystems where water movement can significantly impact detection probabilities.

The observed differences in species composition between surface water eDNA sampling and bottom trawling emphasize the impact of methodological choices on biodiversity assessments (Aubry et al., 2024). It is arguable that relying exclusively on either method may under-represent species diversity, indicating a complementary perspective. Surface water sampling was chosen for its ease and efficiency, making it a non-invasive, more cost-effective and much less time-demanding option compared to bottom trawling. However, its limitations in detecting benthic species suggest that including eDNA samples from bottom water layers could improve detection of demersal taxa and help bridge some of the methodological gaps observed. While this study did not incorporate bottom-layer eDNA sampling, the suggestion to include it reflects the broader aim of assembling a toolkit that captures a more comprehensive view of estuarine biodiversity. Overall, the significant differences in species composition between surface water eDNA metabarcoding and bottom trawling highlight the methodological scope, species detection capabilities, and inherent biases of each approach.

### Beyond the net: eDNA edge in estuarine ecological assessment

The integration of eDNA metabarcoding within the AFI framework has revealed distinct differences in ecological status compared to traditional bottom trawl-based assessments. As previously discussed, these differences are primarily driven by variations in the species assemblages captured by each method, which subsequently influence the index computation. This influence has contributed to the lack of agreement between eDNA- and trawl-derived AFI scores, as indicated by Cohen’s kappa values, and suggests that the two methods assess ecological status based on different species assemblages rather than providing directly comparable results.

The RCR analysis provides further insight into how different species groups influenced overall AFI scores across the two methods. In bottom trawl-derived AFI assessments, resident species (43.7%) had the strongest influence, largely driven by crustaceans. The relevance of crustaceans in the bottom trawl-based AFI was evident that there was poor agreement when bottom trawl AFI calculations without crustaceans, underscoring their critical role, particularly as key component of the AFI for the ecological assessment of some Basque estuaries types (Borja et al., 2004). In contrast, crustaceans were not represented in our eDNA results due to the use of fish-specific primers (teleo region from the 12S gene; Valentini et al. (2016)), which were selected to target vertebrates. AFI scores. Instead, eDNA-derived AFI scores were primarily influenced by omnivorous species (56.7%), reflecting eDNA’s broader taxonomic detection range and its ability to detect species from multiple trophic levels by capturing genetic material dispersed in the water column. These findings underscore the necessity of calibrating eDNA-based assessments to account for taxonomic exclusions and differences in community representation.

A major challenge in integrating eDNA-based ecological assessments is the lack of specific reference conditions. Since the AFI was originally developed using bottom trawl-derived data, its classification thresholds and ecological quality class boundaries are calibrated for datasets dominated by demersal and benthic species. However, eDNA metabarcoding detects species that may not be well represented in bottom trawl surveys. Consequently, applying trawl-derived reference conditions to eDNA-based species lists may introduce classification inconsistencies and differences in ecological assessments. Establishing eDNA-specific reference conditions is therefore essential to ensure that eDNA-based assessments produce ecologically meaningful and comparable results. Once specific eDNA-based reference conditions are implemented, greater convergence between eDNA- and bottom trawl-derived ecological status is expected.

Beyond reference conditions, the integration of eDNA into AFI assessments also requires careful intercalibration with traditional methods (Lepage et al., 2016). Since trawl-based AFI has already been calibrated with other morphological fish sampling techniques under European regulatory frameworks (European Commission, 2024), extending this process to eDNA metabarcoding would help refine indicator species weightings, detection thresholds, and index calculations. Such adjustments are essential for ensuring regulatory compatibility and methodological consistency across monitoring programs. Despite the challenges discussed in the previous section, eDNA metabarcoding presents a compelling alternative in scenarios where conventional capture techniques are impractical or ecologically disruptive. Its non-invasive nature minimizes habitat disturbance while offering a scalable and cost-effective tool for estuarine ecological assessments.

### Way forward

This study highlights the transformative potential of eDNA metabarcoding for estuarine ecological assessment, but strategic advances are needed to ensure its effective integration and long-term applicability. First, the development of eDNA-specific reference conditions is essential to ensure that ecological status classifications are comparable to traditional AFI assessments. This requires the development of intercalibration protocols, recalibration of quality class boundaries and refinement of detection thresholds to improve compatibility and consistency of ecological assessments. Second, addressing taxonomic gaps in genetic reference databases remains a priority to minimize false negatives and misclassifications. Additionally, optimizing sampling strategies by incorporating bottom water samples could improve the detection of demersal taxa, underrepresented in eDNA samples collected from the water surface. Moving forward, we recommend the development of novel eDNA-specific indices tailored to molecular data characteristics, which will enhance their application in estuarine ecological assessment. With these advancements, eDNA metabarcoding can evolve into a standardized, non-invasive tool for long-term estuarine monitoring and assessment.

## Acknowledgements

This investigation was funded by the Basque Water Agency (URA) through a convention with AZTI, by the Department of Economic Development and Infrastructure of Basque Government (project GENGES), by the Spanish Ministry of Science and Innovation (project EDAMAME; reference CTM2017-89500-R) and by the European Union’s Horizon Europe program (projects GES4SEAS with grant agreement no. 101059877 and OBAMA-NEXT with grant agreement no. 101081642). This work was conducted under a doctoral grant awarded to Mukesh Bhendarkar by the Indian Council of Agricultural Research (ICAR), Government of India.

