## Supplementary figures and images for "Advancing ecological assessment: The integration of eDNA metabarcoding into an estuarine fish index"

### Figure S1

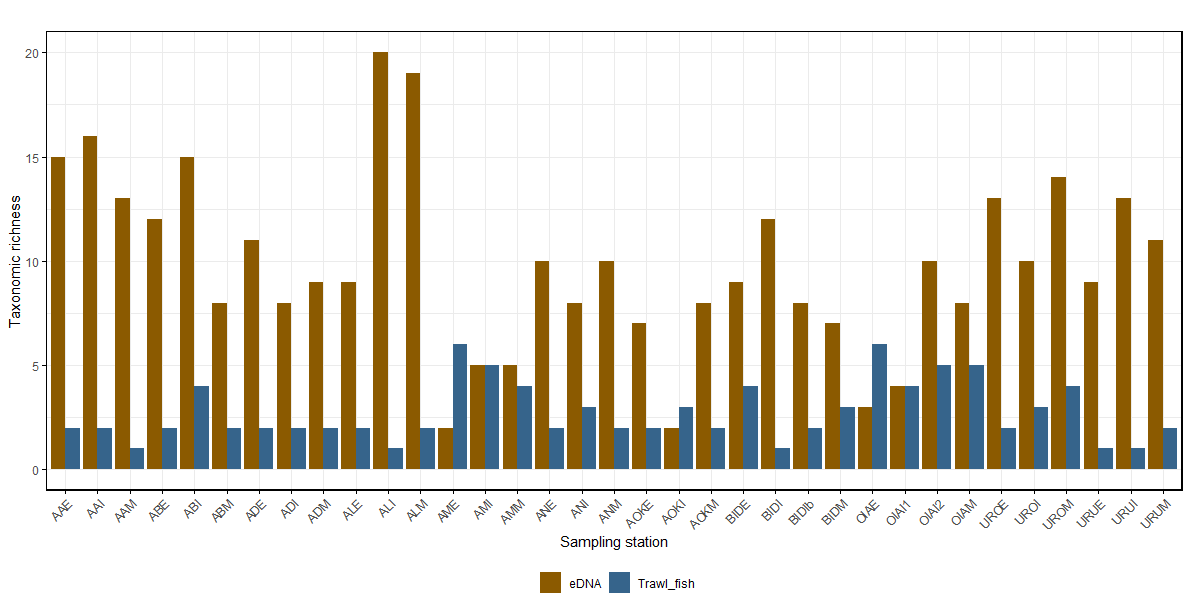

Fig S1. Taxonomic richness between eDNA metabarcoding and trawling survey
